# Itaconic acid production by co-feeding of *Ustilago maydis*: a combined approach of experimental data, design of experiments and metabolic modeling

**DOI:** 10.1101/2023.10.13.562154

**Authors:** A.L. Ziegler, L. Ullmann, M. Boßmann, K.L. Stein, U.W. Liebal, A. Mitsos, L.M. Blank

## Abstract

Itaconic acid is a platform chemical with a range of applications in polymer synthesis and is also discussed for biofuel production. While produced in industry from glucose or sucrose, co-feeding of glucose and acetate was recently discussed to increase itaconic acid production by the smut fungus *Ustilago maydis*. In this study, we investigate the optimal co-feeding conditions by interlocking experimental and computational methods. Flux balance analysis indicates that acetate improves the itaconic acid yield up to a share of 40 % acetate on a carbon molar basis. A design of experiment results in the maximum yield of 0.14 itaconic acid per carbon source from 100 g L^−1^ glucose and 12 g L^−1^ acetate. The yield is improved by around 22 % when compared to feeding of glucose as sole carbon source. To further improve the yield, gene deletion targets are discussed that were identified using the metabolic optimization tool OptKnock. The study contributes ideas to reduce land use for biotechnology, by incorporating acetate as co-substrate, a C2-carbon source that is potentially derived from carbon dioxide.

## 1 INTRODUCTION

The plant pathogen *Ustilago maydis* is a well-established model organism in the field of cell biology, and is in the last decade used in biotechnology due to its natural production of a wide range of value-added molecules [1, 2, 3]. The highly versatile product spectrum contains organic acids such as itaconic, malic, and succinic acid, polyols, and extracellular glycolipids. These molecules are considered value-added chemicals with potential applications in the pharmaceutical, food, and chemical industries [4, 5, 6]. Via metabolic engineering, this broad spectrum was narrowed down specifically to itaconic acid [7]. Itaconic acid itself is a bio-derived platform chemical with uses ranging from polymer synthesis to biofuel production [8].

Currently, industrial itaconic acid production is performed using the filamentous fungus *Aspergillus terreus* [9, 10]. However, itaconic acid production with *A. terreus* displays a challenging process as its production depends on a filamentous morphology, which is required for its high productivity, leading to an increase in production costs [11, 12, 7]. In contrast, advantages of itaconic acid production using *U. maydis* are its unicellular yeast-like growth, high productivity, yields, and titers with a reduced byproduct spectrum due to metabolic engineering [6, 13, 14]. By combining metabolic engineering strategies and an integrated process design, a maximum itaconic acid titer of 220 g L^−1^ was achieved, while for the last 70 years, itaconic acid has been produced using *A. terreus*, reaching titers of 130 g L^−1^ [11]. The current biotechnological processes for itaconic acid production with *A. terreus* and *U. maydis* are based on carbohydrates, such as molasses and glucose [10, 15]. With the overarching goal of a carbon-neutral itaconic acid production process, Ullmann et al. [15] investigated the potential to (co-)consume CO_2_-derived C2 compounds such as acetate.

Since *U. maydis* is a model organism, efficient genetic tools are available, such as CRISPR/Cas [16], high-quality draft genome sequences as well as annotated genomes [17, 18]. Further, a genome-scale metabolic model (GSMM) for *U. maydis* iUma22 was published by Liebal et al. [19]. Those tools enable the investigation of pathogenic mechanisms, biodiversity within the Ustilaginaceae family and biotechnological applications like the metabolic engineering of secondary metabolite production of *U. maydis*.

With the genome-scale metabolic model as a basis, flux balance analysis (FBA) gives insights into the metabolism of the organism [20, 21]. To further enhance the product yield, several bilevel optimization formulations exist to suggest optimal modifications [22, 23, 24, 25, 26]. By deleting a gene, the reactions corresponding to this gene cannot carry flux anymore and hence, the production of, e.g., unwanted side products can be eliminated. The formulation OptKnock by Burgard et al. [22] is a simple, but powerful algorithm to suggest optimal modifications of the metabolic network.

A recent study by Park et al. [27] exploited the potential of co-feeding strategies, boosting bioproduct synthesis. Since each substrate has unique efficiencies for carbon, energy and cofactor generation, varying the relative amounts of substrates in the mixture allows fine-tuning of carbon to-energy-to-cofactor ratios [27]. CO_2_-based substrates such as acetate and formate as carbon sources recently gained interest and show potential for improved itaconic acid production with various Ustilaginaceae. Nevertheless, their utilization remains challenging. Acetate at higher concentrations is toxic, whereas co-utilization with glucose challenges the underlying regulatory network of microbes favoring sugar utilization [28, 29, 30, 31].

Herein, we investigate the co-substrate metabolization of acetate by *U. maydis* MB215 focusing on DoE cultivation experiments concerning growth rate, substrate consumption rates and itaconic acid production. We compare experimental results to FBA results to gather further insights into *Ustilago’s* metabolism. The DoE experiments resulted in growth and substrate uptake data to improve predictions of the GSMM based FBA simulations. We conduct FBA, resulting in gene knockout predictions via OptKnock identifying three knockouts that could potentially enhance itaconic acid production.

## 2 METHODS

### 2.1 Culture conditions

For the cultivation experiments, *Ustilago maydis* strain MB215 was used. Growth and production experiments were performed using modified Tabuchi medium (MTM) according to Geiser et al. [32] containing 0.2 g L^−1^ MgSO_4_·7H_2_O, 0.01 g L^−1^ FeSO_4_·7H_2_O, 0.5 g L^−1^ KH_2_PO_4_, 1 mL L^-1^ vitamin solution, 1 mL L^-1^ trace element solution, and as buffer 19.5 g L^−1^ 2-(N-morpholino) ethanesulfonic acid (MES) or 33 g L^−1 -1^ CaCO_3_ [32]. The carbon sources glucose and sodium acetate were used at changing ratios, see text for detail. NH_4_Cl was added in indicated concentrations. The vitamin solution contained (per liter) 0.05 g D-biotin, 1 g D-calcium pantothenate, 1 g nicotinic acid, 25 g myo-inositol, 1 g thiamine hydrochloride, 1 g pyridoxol hydrochloride, and 0.2 g para-aminobenzoic acid. The trace element solution contained (per liter) 1.5 g EDTA, 0.45 g ZnSO_4_·7H_2_O, 0.10 g MnCl_2_·4H_2_O, 0.03 g CoCl_2_·6H_2_O, 0.03 g CuSO_2_·5H_2_O, 0.04 g Na_2_MoO_4_·2H_2_O, 0.45 g CaCl_2_·2H_2_O, 0.3 g FeSO_4_·7H_2_O, 0.10 g H_3_BO_2_, and 0.01 g KI. Cultivation experiments were performed at 30 °C.

The Design of Experiment (DoE) was performed in System Duetz® (24 deep-well microtiter plates, EnzyScreen, Heemstede, The Netherlands) with a filling volume of 1.5 mL (300 rpm, 80 % humidity, d = 50 mm, Infors HT Multitron Pro shaker, Bottmingen, Switzerland) [33]. Cultures were inoculated in parallel into multiple microtiter plates to a final OD_600_ of 0.5 with cells from an overnight culture in MTM. Tested media combinations are listed in the Appendix (Table 1 and 2). A complete plate was taken as a sacrificial sample for each sample point to ensure continuous oxygenation. Samples for analytical methods were taken at 8 time points distributed throughout the experiment. Experiments were terminated latest after 168 - 240 h when a stable itaconic acid production was observed.

**TABLE 1.** Experimental conditions for first Design of Experiments approach. Conditions to be tested were generated with the software Design Expert. Cultivations were performed in System Duetz® 24 deep-well microtiter plates at 30 °C using MTM with MES buffer and 4 g L^−1^ NH_4_Cl.

| Condition no. | Replicates | Glucose [g L<sup>-1</sup>] | Acetate [g L<sup>-1</sup>] |
| --- | --- | --- | --- |
| 1 | 5 | 9.6 | 6.25 |
| 2 | 5 | 20.0 | 2.5 |
| 3 | 5 | 20.0 | 10.0 |
| 4 | 5 | 45.0 | 0.9 |
| 5 | 8 | 45.0 | 6.25 |
| 6 | 5 | 45.0 | 11.5 |
| 7 | 5 | 70.0 | 2.5 |
| 8 | 5 | 70.0 | 10 |
| 9 | 5 | 80.3 | 6.25 |

**TABLE 2.** Experimental conditions for second Design of Experiments approach. Conditions to be tested were generated with the software Design Expert. Cultivations were performed in System Duetz® 24 deep-well microtiter plates at 30 °C using MTM with CaCO_3_ buffer and 0.8 g L^−1^ NH_4_Cl.

| Condition no. | Replicates | Glucose [g L<sup>-1</sup>] | Acetate [g L<sup>-1</sup>] |
| --- | --- | --- | --- |
| 1 | 2 | 26 | 7 |
| 2 | 3 | 30 | 2 |
| 3 | 3 | 30 | 12 |
| 4 | 2 | 65 | 1.4 |
| 5 | 8 | 65 | 7 |
| 6 | 2 | 65 | 12.5 |
| 7 | 3 | 100 | 2 |
| 8 | 3 | 100 | 12 |
| 9 | 2 | 104 | 7 |

| Sample ID | Number of samples | Average growth rate [h<sup>-1</sup>] | Variance |
| --- | --- | --- | --- |
| 1 | 5 | 0.0464 | $6.23 \times 10^{-5}$ |
| 2 | 5 | 0.0391 | $1.83 \times 10^{-2}$ |
| 3 | 5 | 0.0519 | $1.56 \times 10^{-2}$ |
| 4 | 5 | 0.0472 | $9.13 \times 10^{-6}$ |
| 5 | 8 | 0.0223 | $1.76 \times 10^{-5}$ |
| 6 | 5 | 0.0105 | $9.91 \times 10^{-4}$ |
| 7 | 5 | 0.0225 | $1.28 \times 10^{-5}$ |
| 8 | 5 | 0.0076 | $4.22 \times 10^{-4}$ |
| 9 | 5 | 0.0223 | $1.09 \times 10^{-4}$ |

### 2.2 Process Analytics

Cell growth was determined by measuring the optical density at 600 nm (OD_600_) with an Ultrospec 10 Cell Density Meter (Amersham Biosciences). Carbon sources and metabolites such as glucose, acetate, itaconate, malate, succinate and erythritol in the supernatant were analyzed via high-performance liquid chromatography (HPLC). All samples were filtered with Rotilabo® syringe filters (CA, 0.20 *μ*m) and were afterward diluted in a range of 1:5 – 1:50 with 5 mM H_2_SO_4_. Supernatants were analyzed in a DIONEX UltiMate 3000 HPLC System (Thermo Scientific) with a Metab-AAC column (300 x 7.8 mm column, ISERA). Elution was performed with 5 mM H_2_SO_4_ at a flow rate of 0.6 mL min^-1^, and a temperature of 40 °C. For detection, a SHODEX RI-101 detector (Showa Denko Europe GmbH) and a DIONEX UltiMate 3000 Variable Wavelength Detector set to 210 nm were used. Analytes were identified via retention time and UV/RI quotient compared to standards. All values are the arithmetic mean of two to four biological replicates. Error bars indicate the standard deviation from the mean. Statistical analysis of significant difference (P-value) was performed using unequal variances t-test with unilateral distribution (P values: < 0.05 =*).

### 2.3 Design of experiments

The software Design Expert 11 (Stat-Ease, Minneapolis, MN, USA) was used to set up and evaluate the Design of Experiments (DoE). A response-surface, 3-factor central composite design (CCD) was chosen to simultaneously evaluate the influence of varying concentrations of glucose and sodium acetate on the growth of *U. maydis* MB215 as well as the itaconic acid production. For the DoE predicting the effect of co-feeding glucose and acetate on the growth rate, nine different conditions were tested. All conditions were performed as 4-fold replicates, except the central condition (45 g L^−1^ glucose and 6.25 g L^−1^ acetate) as 8-fold replicates (Table 1, Appendix). The model predicting itaconic acid production during acetate co-metabolization cultivations was set up as stated in the following. Nine different conditions were tested, whereas the approaches with the lowest (26 g L^−1^) and highest (104 g L^−1^) glucose concentration, as well as the approaches with the lowest and highest acetate concentration, were carried out as duplicates. All remaining medium-level (30 - 100 g L^−1^) glucose conditions were implemented in triplicates, except the central condition (65 g L^−1^ glucose and 7 g L^−1^ sodium acetate), as an octuplet (Table 2, Appendix). To enhance the significance of the model, each condition was tested in replicates. Factors showing a significant influence (p-value <0.05) were incorporated into the model. If a factor with a non-significant influence was substantial for maintaining the model hierarchy, exceptions were made. The input data were transformed, if necessary, according to the recommendations of the software. To prove the predicted conditions and confirm the design, validation in 24-deep well plates and up-scaling shaking flasks were performed.

### 2.4 Flux balance analysis and genome-scale model

FBA determines the reaction fluxes within an organism when the metabolism is at steady state [20, 21], and is based on the metabolic network. The metabolic network includes all known reactions and metabolites inside the cell, together with the stoichiometric factors of the reactions and information on the enzymes catalyzing the reactions. The reaction fluxes are determined by solving the following optimization problem, a linear program:

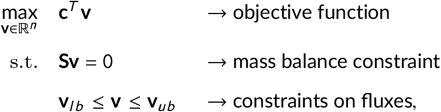

where **v** denotes the flux vector with *n* entries, **S** is the stoichiometric matrix, and **v**_*l b*_ and **v**_*ub*_ are parameter vectors imposing lower and upper bounds on the fluxes, respectively. The objective function represents, e.g., the biomass production or the target chemical production, by selecting appropriate values for the cost vector **c**^*T*^ . The mass balance constraint represents mass conservation for each metabolite, assuming steady state.

The FBA computations are based on an updated version (v1.1) of the genome-scale metabolic model of *U. maydis* [19]. In model v1.1, the exchange reaction for formate was added, and CO_2_ uptake was enabled up to a rate of 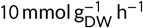. To prevent non-physical reactions fluxes, in model v1.1, the following reactions were defined as irreversible: acetylphosphate *fructose-6-phosphate phosphoketolase* (E.C. 4.1.2.22), *phosphoketolase* (E.C. 4.1.2.9) and the phosphoenolpyruvate carboxy kinase related reaction *RXN-12481* (E.C. 4.1.1.32). The activity of following reactions resulted in unrealistic overall flux distributions and thus, they were deactivated in v1.1: methanol dehydrogenase *RXN-11453* (E.C. 1.2.98.4), pseudo-catalase *PRDXl* (E.C. 1.11.1.6), and ATP-citrate-proline lyase *ACITL* (E.C. 2.3.3.8). Newly added reactions based on experimental evidence include the mannitol production from mannose-6-P via the pseudo reaction *PseudoMan6P2Mnl* [34] and cytoplasmic isocitrate lyase activity *ICL*. The updated model is available on the *U. maydis* GSMM Github page as v1.1 (https://github.com/iAMB-RWTH-Aachen/Ustilago_maydis-GEM/tree/v1.1).

For investigating co-feeding conditions, the uptake of glucose and acetate were varied. To keep results comparable, the total flux of carbon atom was kept constant. Glucose has six carbon atoms and acetate has two carbon atoms, and hence, the equation

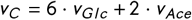

was set as additional constraint. When setting the total carbon flux and varying the acetate flux, the corresponding glucose flux was calculated during FBA. using the software COBRAPy [35] and the open source solver GLPK. The code for the FBA computations is openly available (https://github.com/iAMB-RWTH-Aachen/Ustilago_maydis-GEM/tree/master/data/AcetateCofeed).

### 2.5 Optimal genetic modifications

Genetic modifications influence the flux through a metabolic network. Deletion of a certain gene inhibits the flux through the reaction associated to this gene. Optimal reaction knockouts were suggested by using the bilevel optimization formulation OptKnock [22]:

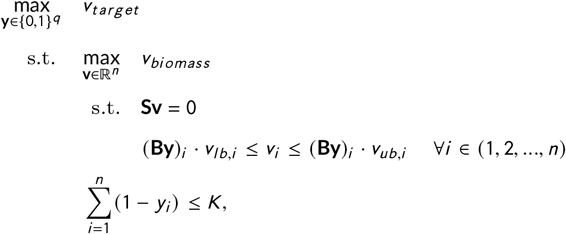

where the vector **y** denotes the knockouts, i.e., *y*_*i*_ = 0 indicates knockout of reaction *i* . The maximum number of knockouts is described by *K* . Reversible reactions are transformed into two irreversible reactions. The original number of—possibly reversible—reactions is denoted with *q*, the number of reactions in the irreversible network is *n*. The matrix **B** is a matrix that maps the irreversible reactions to their original—possibly reversible—reaction. For creating the irreversible network, the software COBRAtoolbox [36] was used. The upper level of the optimization program describes the bioengineering objective to maximize the flux of the target product, whereas the lower level describes the microorganism that aims to maximize the biomass flux. The reformulated form of OptKnock as a single-level program was implemented in the optimization language libALE [37] and solved using libDIPS [38] with gurobi 9.5.2 [39] as solver. The code for the OptKnock computations is openly available (https://github.com/iAMB-RWTH-Aachen/Ustilago_maydis-GEM/tree/master/data/AcetateCofeed).

## 3 RESULTS AND DISCUSSION

With *Ustilago maydis*, itaconic acid is produced in a two-phase process. An initial growth phase is followed by the production phase upon depletion of a growth-limiting nutrient, here, itaconic acid production is induced under nitrogen limitation [40]. Thus, the total amount of itaconic acid produced in a process is also dependent on the amount of biomass produced. If the growth-limiting nutrient is provided in a higher concentration, more biomass will be formed. This will likely increase the volumetric productivity, but will also decrease the overall yield, since more carbon source is consumed in the formation of the biomass. The trade-off of yield and rate under co-feeding conditions was further examined by simulations and wet-lab experiments.

### 3.1 Comparing the experimental growth rate with the predicted growth rate under co-feeding conditions

First, we evaluated the growth rate under nine different co-feeding conditions (Appendix, Table 1). The results corroborate previous findings that higher glucose concentrations result in lower growth rates [19] and this sensitivity is increased by higher acetate concentrations. The co-feeding with 10 g L^−1^ acetate and 20 g L^−1^ glucose (#3 with ≈17 % relative acetate C-mol ratio in Figure 1a) resulted in the highest observed growth rate of 0.066 h^−1^.

**FIGURE 1.**
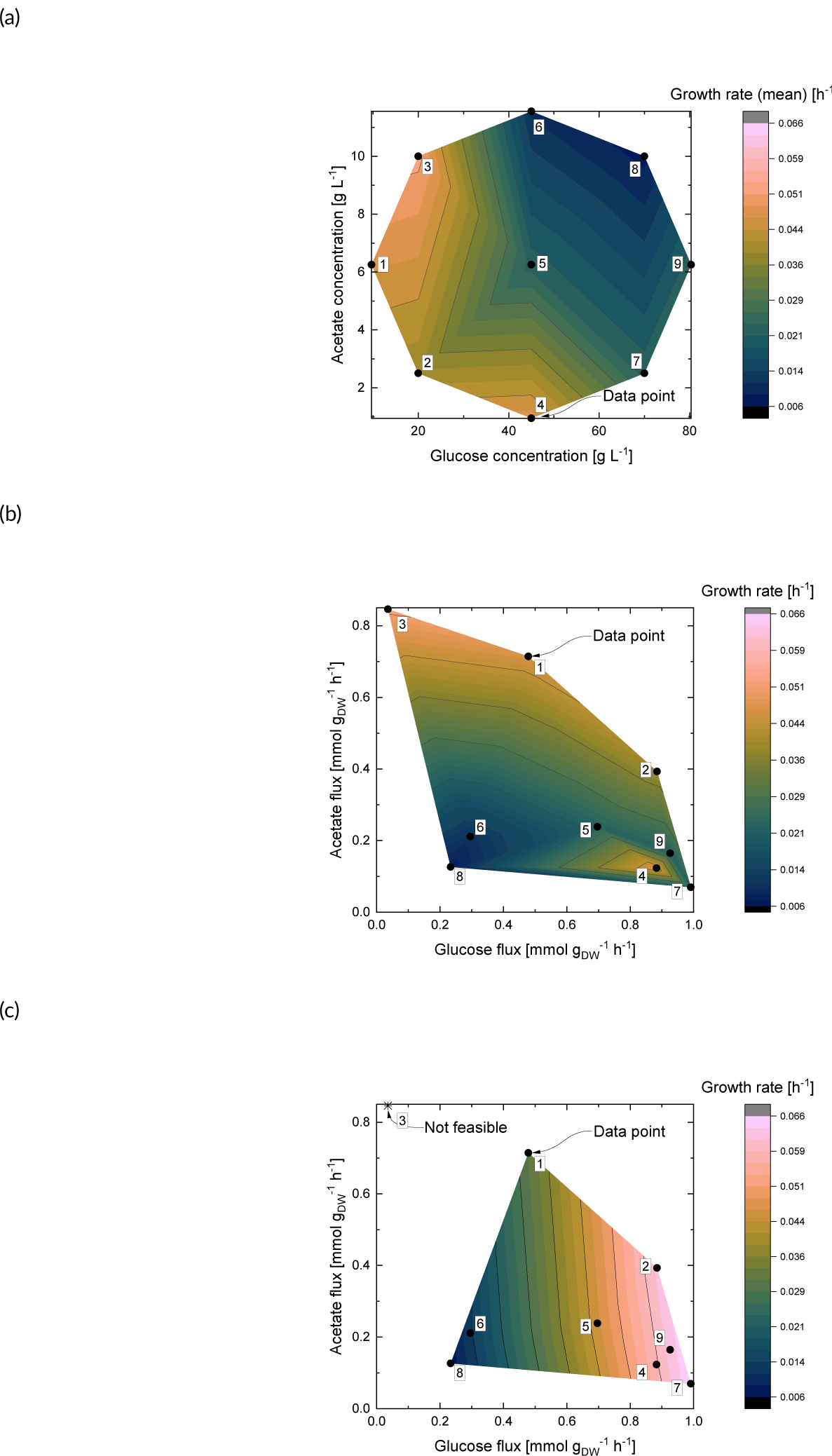
Growth rate over glucose and acetate flux. Subfigure (a) shows experimental data on growth rate over glucose and acetate concentration obtained during the first DoE cultivation experiment using *U. maydis* MB215. System Duetz® 24-deep-well plate cultivations were performed with MTM containing MES buffer and 4 g L^−1^ NH_4_Cl at 30 °C. Growth rate is averaged. Subfigure (b) shows experimental data with nine data points, data points are arithmetic means from 5 to 8 measurements. Subfigure (c) shows the theoretical growth rate over glucose and acetate flux predicted by FBA. Nine points are simulated of which one is not feasible.

To get further insights into how the metabolic configuration changes with respect to the co-feeding conditions, we measured the uptake rates that correspond to the nine co-feeding conditions (Figure 1b). There are three sets of measurements with increasing glucose and constant acetate concentrations (sets #2+#7, #1+#5+#9, #3+#8). All sets display a pronounced decrease in acetate uptake rate with a modest increase in glucose uptake rate. For example, for the measurements #1+#9, the glucose concentration increases from 10 to 80 g L^−1^, the glucose uptake rate increases only from ≈ 0.45 to 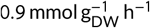, while despite constant level the acetate uptake decreases from ≈ 0.7 to below 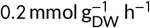. The same effect is exerted by acetate on constant glucose levels: increasing the acetate concentration decreases glucose uptake rates (sets #2+#3, #4+#5+#6, #7+#8). In set #7+#8, whereas the glucose concentration remains constant, the glucose uptake rate decreases from ≈ 1 to 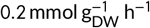 with an acetate concentration increase from 2.5 to 10 g L^−1^. Note the growth anomaly for the measurements #4, #7 and #9. Although the substrate uptake rates for glucose and acetate are comparable for the three measurements, the increase of acetate concentration causes a drop in the yield with conditions #7 and #9. Overall, the concentration growth profile in Figure 1a is reversed in the substrate uptake rate profile in Figure 1b. Measurements with high concentrations are mirrored to low uptake rates (e.g., #8) and vice versa (e.g., #2). Acetate causes weak acid stress and hence, the cell responds with reduced uptake. At high glucose and acetate concentrations in the medium, substrate uptake is reduced. Maassen et al. [40] showed that lower concentrations of glucose are likely to reduce osmotic stress, which has a positive effect on cellular growth.

The inhibiting effect of acetate on glucose consumption is absent in iUma22 for growth optimization. When comparing the experimental with the simulated growth rate in Figure 1c, the growth rates are overall in the same range. Especially, we find equivalent growth rates at low glucose and acetate fluxes (measurements #8 and #6). The results divert with increasing glucose and acetate flux. At conditions with high acetate flux, the experimental growth rate exceeds the simulated growth rate. One example for this is point #1, where the experimental growth rate is 0.046 h^−1^, whereas the simulated growth rate is 0.03 h^−1^. This low simulated growth rate is no outlier but following the trend lines of the simulation. The co-feeding conditions at point #3 do not furnish a feasible result, which matches the observations in Section 3.3.

### 3.2 Itaconic acid production: Design of experiments and metabolic modeling enable finding the optimal co-feeding strategy

In a second DoE, for which we hypothesized combinatorial effects of cell growth, substrate uptake rates as well as itaconic acid production, nine different media combinations were tested (Appendix, Table 2). The DoE study returned the equation

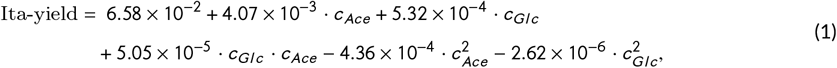

for the calculation of itaconic acid yield from co-feeding conditions, where *c*_*Gl c*_ and *c*_*Ace*_ are the concentrations of glucose and acetate, respectively. The equation uses the practically relevant substrate concentration as variable, and for comparison with the FBA a transformation of the concentrations into fluxes is necessary. For the transformation, we conducted an experimental study to correlate starting concentrations and uptake rates of glucose and acetate. The following linear regression models result in a coefficient of determination (*R* ^2^) 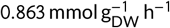 for *v*_*Gl c*_ and 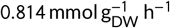 for *v*_*Ace*_ :

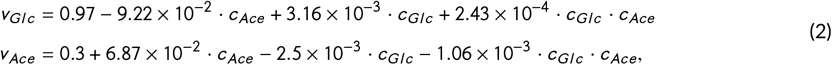

where *v* are fluxes, *c* are the concentrations and *Gl c* and *Ace* represent glucose and acetate, respectively. Note, that the glucose uptake flux is strongly and inversely controlled by the acetate concentration. The additional interaction term *c*_*Gl c*_ · *c*_*Ace*_ displays the interactions between glucose and acetate concentrations. The correlation enables us to display the yield of itaconic acid over the glucose and acetate flux, which makes it comparable to the results from the FBA. In the FBA, we set a threshold on biomass, which is 50 % of the maximal theoretical biomass flux at this point. Doing so, the FBA furnished results that were closer to experimental observations, as the organism uses the carbon substrate for product formation as well as biomass formation. Figure 2 shows the results from the design of experiments study in Subfigure 2a and the results from the FBA in Subfigure 2b.

**FIGURE 2.**
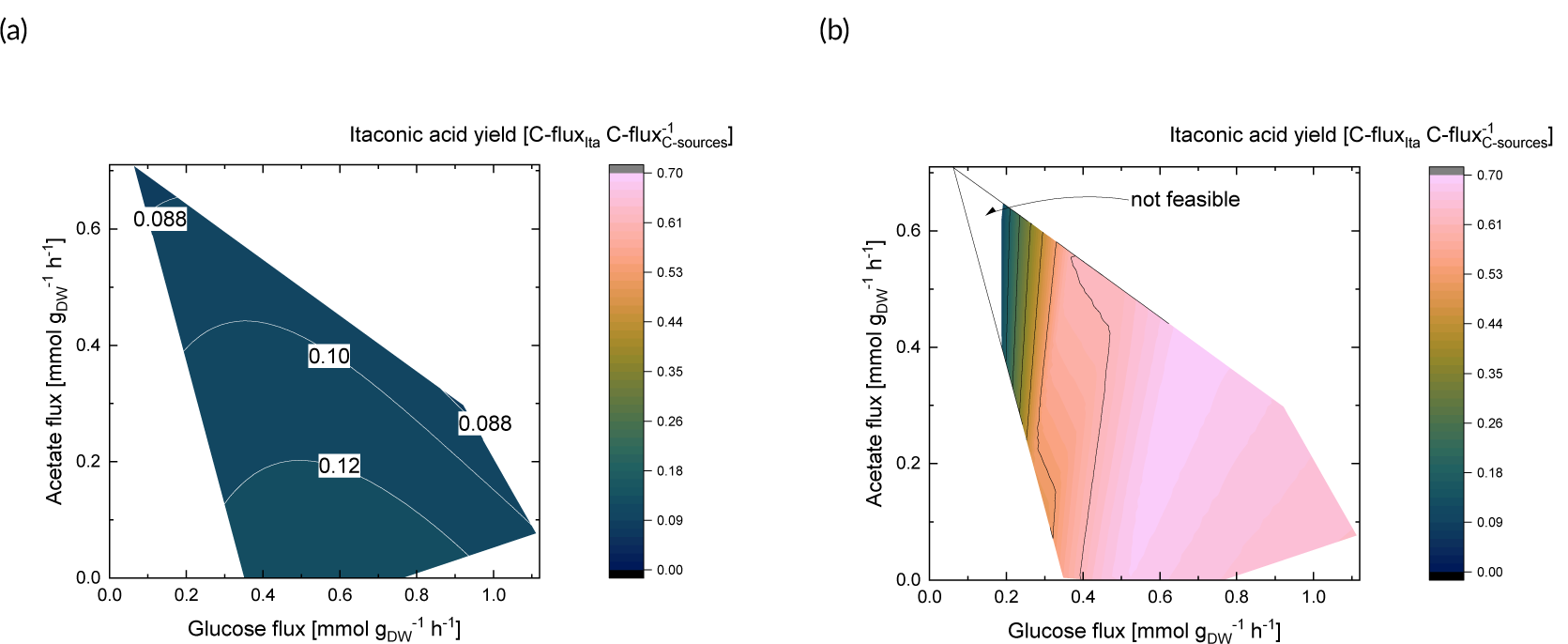
Itaconic acid yield depending on glucose and acetate uptake. The lines correspond to a constant yield. Subfigure (a) shows results from the design of experiments study of itaconic acid yield over acetate and glucose flux. Glucose flux and acetate flux were calculated from acetate and glucose concentrations via correlation in Equation 2, and itaconic acid yield was calculated via Equation 1. System Duetz® 24-deep-well plate cultivations were performed with MTM containing CaCO_3_buffer and 0.8 g L^−1^ NH_4_Cl at 30 °C. Subfigure (b) shows FBA results of itaconic acid yield over acetate and glucose flux. Itaconic acid yield was calculated via FBA, with a threshold on biomass growth rate of 50% of the maximal theoretical biomass growth at each point.

The corresponding glucose concentration ranges from 26 g L^−1^ to 80 g L^−1^, whereas the acetate concentration ranges from 1.5 g L^−1^ to 11.5 g L^−1^. In Figure 2a, we observe the level of itaconic acid yield as a plateau without steep descends. More precisely, the yield of itaconic acid is 0.09 to 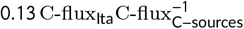. The lowest yield corresponds to the highest acetate flux, whereas the highest yield is achieved when the acetate flux is low. From the FBA displayed in Figure 2b, we find that below a glucose flux level of around 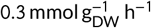, there is a descent. Below a glucose flux of around 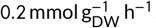, there is no feasible solution, which is in coherence with the observations in Section 3.3. Above a glucose flux of 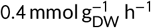, we find a plateau. The yield is predicted in the range of 0.12 to 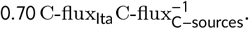. The lowest yield corresponds to the lowest glucose flux, whereas the highest yield is observed in a medium region of around 0.5 to 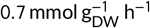 glucose. Notably, the predicted yield of itaconic acid is higher than the experimental yield. The experimental yield is decreased by the formation of by-products *in-vivo*, unaccounted by the model. The results of the design of experiments analysis indicate that the itaconic acid yield is mainly dependent on the acetate uptake. In contrast, in the FBA, the glucose uptake appears to be decisive. One explanation for this phenomenon can be the toxicity of acetate to the microorganism. This toxicity is not included in FBA. Lastly, we observe a plateau with both methods, which indicates a coherence.

During DoE cultivation experiments with *U. maydis* MB215, the itaconic acid yield increased with increasing glucose concentration and for increasing acetate concentration when considering the equally high glucose concentrations. The maximum yield was reached with 0.14 ± 0.00 C-mol_Ita_/C-mol_C-source_ for the condition with the highest total amount of carbon with 100 g L^−1^ glucose and 12 g L^−1^ acetate. To get insights on the influence of glucose and acetate co-feeding on the itaconic acid yield compared to a standard glucose feed, cultivations with similar C-mol were performed for both (Figure 3).

**FIGURE 3.**
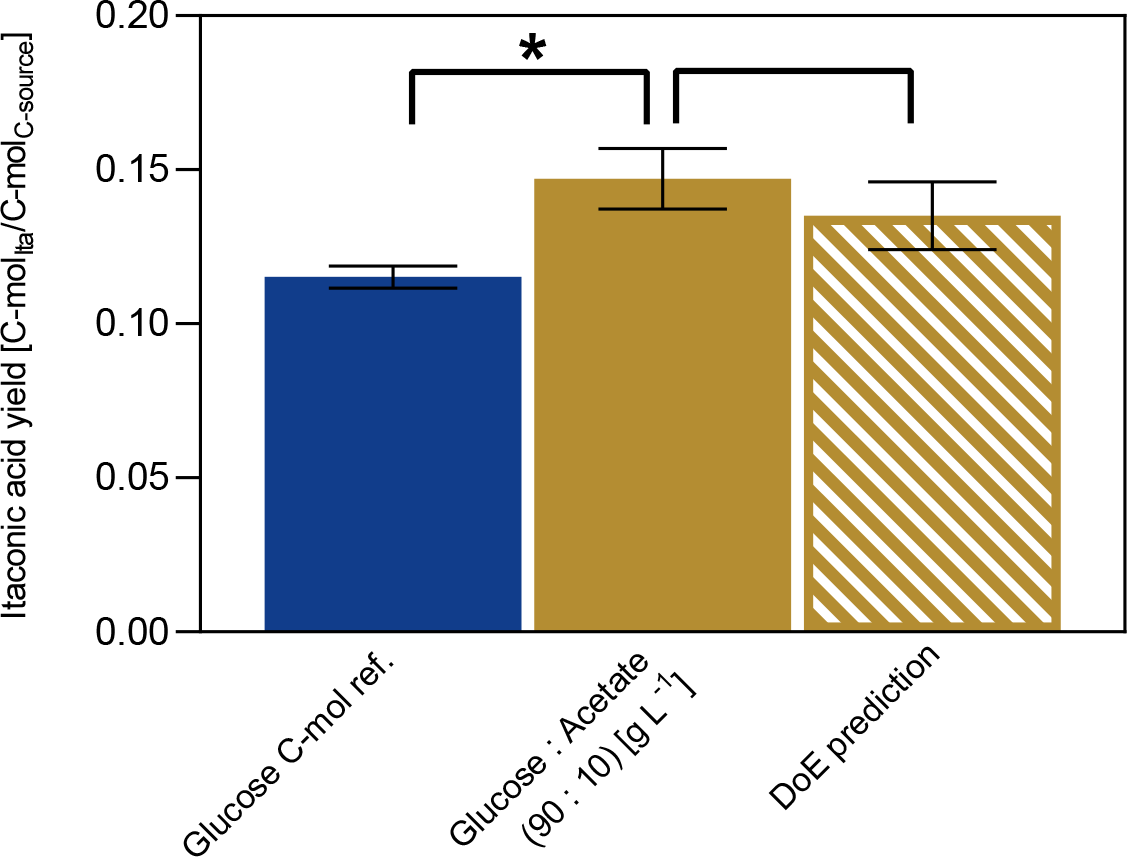
Comparison of itaconic acid yields for glucose and acetate co-feeding with standard glucose cultivation. The co-feeding of glucose and acetate (brown) was compared to the DoE prediction (shaded brown) and a glucose reference containing the same amount of total carbon (blue). System Duetz® 24-deep-well plate cultivations were performed at 30 °C with 0.8 g L^−1^ NH_4_Cl and CaCO_3_ buffer. Error bars indicate the deviation from the mean for n=3. Statistically significant differences in itaconic acid production (0.01 < p < 0.05) are indicated as *.

For the cultivation, 90 g L^−1^ of glucose and 10 g L^−1^ of acetate were used. As these are no conditions used to create the DoE, the predictive power of this was tested at the same time. Firstly, the cultivation with glucose and acetate reached a yield of 0.15 ± 0.01 C-mol_Ita_/C-mol_C-source_. Compared with the DoE prediction 0.14 ± 0.01 C-mol_Ita_/C-mol_C-source_ the values do not differ significantly. The respective glucose reference cultivation reached a yield of 0.11 ± 0.00 C-mol_Ita_/C-mol_C-source_ which is significantly lower compared to cultivation with acetate (Figure 3). Since the reached yield did not differ significantly from the model prediction, the validation was considered as successful, indicating that the model has an accurate predictive power regarding the yield for a defined substrate combination within the model ranges. Moreover, the comparison to a glucose reference with a similar total carbon amount showed that acetate has a positive effect on the itaconic acid production, which was already indicated in previous experiments [15].

### 3.3 Exploring the range of co-feeding using flux balance analysis

To further explore the range of co-feeding, we determined the theoretical growth rate and the theoretical itaconic acid flux up to a relative acetate carbon-flux of 100 %. Figure 4 shows the results.

**FIGURE 4.**
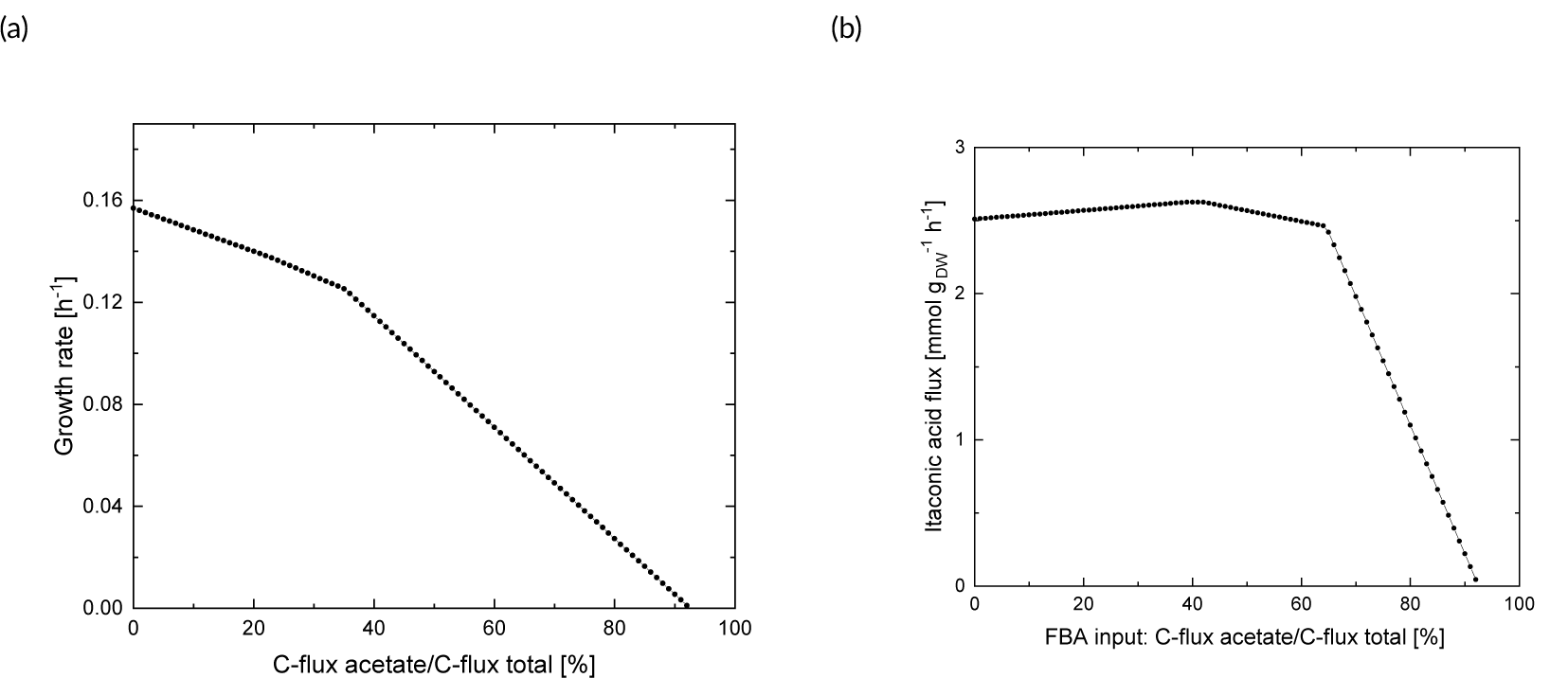
Simulated growth rate and itaconic acid flux over the relative acetate carbon uptake (C-flux) predicted by FBA with total and constant C-mol uptake rate of 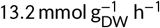. The FBA was optimized for growth in Subfigure (a) and itaconic acid production with no threshold on biomass growth in Subfigure (b).

We conducted the FBA with a carbon flux of 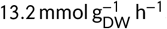, which is equivalent to a glucose uptake rate of 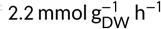 [19] and performed the FBA in steps of 1 %. Although the mechanism of acetate stress is not explicitly included in iUma22, a decrease in the growth rate with increasing relative acetate carbon ratio could be reproduced, as visualized in Figure 4a. Moreover, a bendpoint comes visible at a relative acetate C-flux of 35 %. Interestingly, for ratios of acetate higher than 35 %, there exist several possible solutions for achieving the same optimal growth rate. In the presented solution in Figure 4a, all available carbon is metabolized by the organism. However, other solutions exist where not all available carbon of 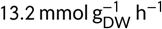 is taken up, and, hence, the metabolic configurations can vary. At around 92 % acetate in the substrate, the solution becomes infeasible. Presumably, not all metabolites included in the biomass exchange reaction, i.e., necessary for biomass production, can be metabolized from acetate.

In order to explore the range of co-feeding for itaconic acid flux, we determined the maximal theoretical itaconic acid flux. Maximal theoretical flux means that we did not impose a threshold on the flux of biomass. The result of this study is shown in Figure 4b. The maximal theoretical flux of itaconic acid on pure glucose (0 % C-mol acetate per C-mol total), is 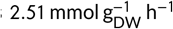, with a yield of around 0.95 mol mol^−1^. Presumably, itaconic acid can only be formed from glucose via losses, for example via the pyurvate dehydrogenase, which also produces CO_2_. The itaconic acid flux rises linearly to a maximum of 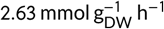 (>0.99 mol mol^−1^ yield), with a corresponding increase in the relative carbon acetate uptake ratio to 41 %. The maximal theoretical flux is reached on a plateau between 41 % and 42 % relative carbon flux acetate. Presumably, one or several metabolites needed for itaconic acid production are synthesized better from acetate than from glucose, which is why a certain share of acetate boosts the itaconic acid production. Further increasing the relative acetate carbon uptake ratio reduces the yield to 0.94 mol mol^−1^ (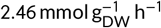 itaconic acid flux) at 64 % relative acetate input. The acetate carbon ratio of 64 % is a threshold, beyond which the overall carbon uptake rate decreases for both substrates and the flux limits are not necessarily reached. Again, this is due to multiple possible solutions inside the metabolic network that fulfil the maximal itaconic acid flux at the respective condition. It comes visible that beyond a share of 92 % carbon flux acetate per carbon flux total, the flux of itaconic acid would become negative, which is not a feasible solution. Presumably, similar to the observations on the simulated growth rate, not all metabolites that are necessary for itaconic acid production, can be metabolized by acetate.

We draw the following conclusions from this study: First, co-feeding of acetate can increase the theoretical flux of itaconic acid in comparison to pure glucose as substrate. Second, the model reproduces the expected growth inhibition on pure acetate. Third, the bend points can indicate that the metabolism switches reaction pathways at these points. Fourth, FBA computations suggest that itaconic acid production on pure acetate is not possible.

To get a better understanding of the flux distributions with varying acetate C-mol ratios in the substrate, the three extreme flux distributions *F1* (corresponding to 1 % C-mol acetate in the substrate), *F2* (41 % C-mol acetate), and *F3* (61 % C-mol acetate) were selected to investigate the metabolic flux details. Figure 5 shows relevant pathways with their normalized carbon fluxes.

**FIGURE 5.**
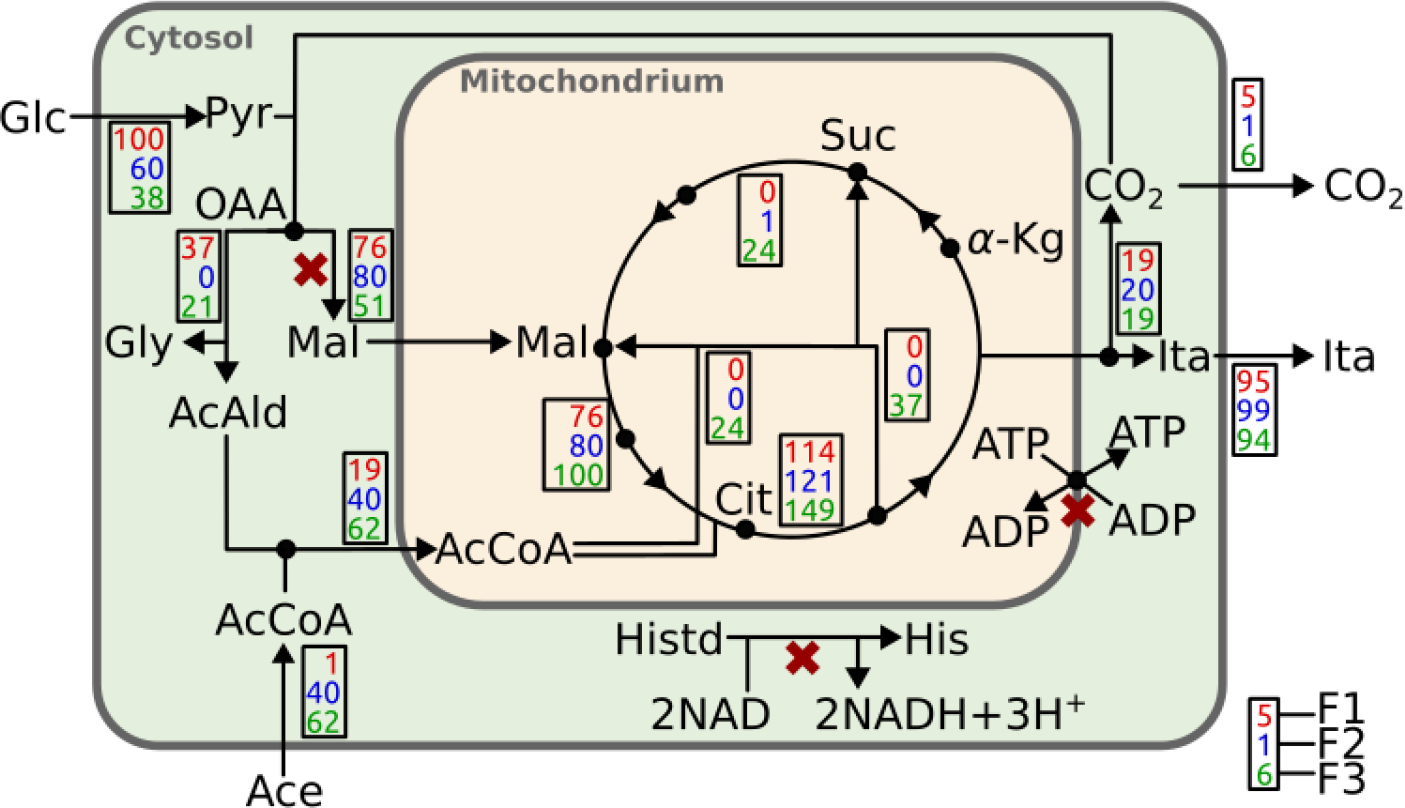
Flux distributions of three simulated flux solutions *F1, F2* and *F3* of Figure 4b for different acetate substrate ratios optimized for itaconic acid production. *F1* corresponds to 1%, *F2* to 41%, and F3 to 61% C-mol acetate in the substrate mix. For comparability, the carbon uptake was normalized to 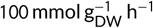 and each reactions shows the conversion rate of participating carbon atoms. The pentose phosphate cycle is inactive, *ALD* represents a pathway including the Threonine Aldolase reaction (E.C. 4.0.2.5) which was used to convert oxaloacetate via threonine to acetaldehyde. The red crosses indicate mutations with increased itaconic acid and biomass production identified by optimization with three allowed knockouts.

Figure 5 shows the strategies of carbon conversion to itaconate for different glucose-acetate uptake ratios. The flux values at each reaction represent the percent of converted carbon atoms, relative to the total carbon atom uptake. For example, the *ALD*-pathway of F1 (red, top number) represents the conversion of 37 oxaloacetate carbon atoms to 18 carbon atoms of glycine (Gly) and 18 carbon atoms of acetaldehyde (AcAld). The pool of acetyl-CoA (AcCoA) is formed both by the 18 AcAld carbons and the uptake of 1 carbon of external acetate (Ace). The resulting 19 carbon atoms of AcCoA relocate into the mitochondria for further reactions in the TCA. The optimal path with highest icaconic acid production (F2, blue) avoids glyoxylate shunt fluxes. While also the glucose only itaconic acid production route displays inactive glyoxylate shunt (F1, red), the AcCoA for the citrate synthase reaction has to be generated from pyruvate. The model chooses a hypothetical route via the threonine aldolase reaction to avoid loosing CO_2_ via pyruvate dehydrogenase. The acetate co-feed avoids the detour route and results in a more efficient glucose carbon utilization.

### 3.4 Improving the itaconic acid yield by knocking out three reactions

Gene knockouts were predicted using OptKnock in a range of one to three knockouts. The carbon input was set to 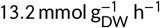 and a threshold on biomass was induced, which was 0.016 h^−1^. Without knockouts, no itaconic acid was produced when conducting a FBA with the objective of biomass formation and 0.16 h^−1^ biomass was formed as only glucose was chosen as carbon source by the optimizer. With one to two knockouts, no significant improvement came visible, meaning that no itaconic acid was formed. We conclude that two knockouts are not sufficient to improve the yield of itaconic acid. With three knockouts, OptKnock predicted a significant improvement in the itaconic acid flux, which was 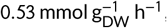, while decreasing the biomass formation to 0.029 h^−1^. Hence, more carbon atoms were used in itaconic acid formation and less in biomass formation, which was intended. Figure 5 shows the reactions that were targeted for knockout by red crosses. The predicted reactions are “TRANS Adenine Nucleotide Transporter”, “Cytosolic MALATE-DEH-RXN” and “HISTD”. The optimizer chose for pure glucose and no acetate as carbon input. When optimizing on glucose from the start, the last reaction of the suggested ones (“HISTD”) changed to “PPCOACm”. This indicates that there are several reactions that lead to the optimal solution. The results from OptKnock, however, cannot guarantee the improved itaconic acid yield. To tackle this drawback, more advanced formulations [23, 26] should be considered in future work. Moreover, “Cytosolic MALATE-DEH-RXN” as well as “TRANS Adenine Nucleotide Transporter” are reactions that actively take part in the itaconic acid production, which is why the knockout of these reactions are questionable with biological experience. Formulations that are suitable for a modified organism [24, 25] should therefore be considered next. Moreover, the suggested knockouts should also be tested experimentally.

## 4 CONCLUSION

FBA suggested that a carbon atom division of 40 % for acetate and 60 % for glucose achieves the highest itaconic acid yield. In the associated optimally simulated pathway, all AcCoA from acetate was used for citrate synthase and each glucose was converted to two malate, further fixing 2 mol CO_2_ per glucose. This additional CO_2_ fixation results into an increased yield on glucose. The mutation simulation targeted high rates whereas the co-feeding flux simulation focused on high itaconic acid yields. The predictions center on glucose related reactions for rate increases whereas the oxalacetic acid to malate reaction is a key reaction for high yields during co-feeding.

Efficient co-feeding rests on a delicate balance between glucose and acetate consumption. The DoE data analysis shows a direct correlation of the itaconic acid yield with the acetate concentrations. However, because the substrates reciprocally inhibit their uptake rates, the rates will deteriorate with increasing concentrations. The simulation indicates a minimum uptake rate of glucose (0.5 mmol/gDW/h) is necessary to profit from acetate. This is corroborated by the DoE, which displays highest acetate and yield robustness at a glucose uptake rate of 0.5 mmol/gDW/h. This fragile substrate balance would have to be overcome for any successful industrial application by increasing robustness via mutations followed by a confirmation of the performance during large-scale cultivations. Also, fed-batch cultivations that keep the carbon source below detrimental concentrations will contribute to harvest the benefits of the CO_2_ derived co-substrate acetate.

## Abbreviations

Ace: acetate
AcCoA: acetyl-CoA
*α*-Kg: *α*-Ketoglutarate
C: carbon
FBA: flux balance analysis
Glc: glucose
Gly: glycine
His: histidine
Histd: histidinol
Ita: itaconic acid
Mal: malate
MES: 2-(N-morpholino) ethanesulfonic acid
MTM: modified Tabuchi medium
Pyr: pyruvate
Suc: succinate
TCA: tricarboxylic acid cycle.

## 5 APPENDIX

### Funding

This project was funded by the Deutsche Forschungsgemeinschaft (DFG, German Research Foundation) under Germany’s Excellence Strategy – Cluster of Excellence 2186 “The Fuel Science Center” – ID: 390919832

### Author contributions

ALZ designed the computations (FBA and OptKnock) and carried out the project management. LU designed the laboratory experiments and the DoE studies. MB performed the computations under the supervision of ALZ. KLS performed the laboratory experiments under the supervision of LU. UWL provided the updated metabolic network. ALZ, LU and UWL analyzed and discussed the data. ALZ, LU, UWL and KLS wrote the manuscript draft. AM and LMB conceptualized the project, discussed the data and reviewed the draft. All authors read and approved the final manuscript.

### Conflict of interest

There are no conflicts to declare.

### Data availability statement

The updated genome-scale metabolic model iUma22 is available in our GitHub repository “Ustilago_maydis-GEM” as v1.1 at https://github.com/iAMB-RWTH-Aachen/Ustilago_maydis-GEM/tree/v1.1. The FBA amd OptKnock computations underlying this article are available in our GitHub repository “Ustilago_maydis-GEM” at https://github.com/iAMB-RWTH-Aachen/Ustilago_maydis-GEM/tree/master/data/AcetateCofeed.

